# Fast-local and slow-global neural ensembles in the mouse brain

**DOI:** 10.1101/2022.07.14.500088

**Authors:** Thomas J Delaney, Cian O’Donnell

## Abstract

Ensembles of neurons are thought to be co-active when participating in brain computations. However, it is unclear what principles determine whether an ensemble remains localised within a single brain region, or spans multiple brain regions. To address this, we analysed electrophysiological neural population data from hundreds of neurons recorded simultaneously across nine brain regions in awake mice. At fast sub-second timescales, spike count correlations between pairs of neurons in the same brain region were stronger than for pairs of neurons spread across different brain regions. In contrast at slower timescales, within- and between-region spike count correlations were similar. Correlations between high-firing-rate neuron pairs showed a stronger dependence on timescale than low-firing-rate neuron pairs. We applied an ensemble detection algorithm to the neural correlation data and found that at fast timescales each ensemble was mostly contained within a single brain region, whereas at slower timescales ensembles spanned multiple brain regions. These results suggest that the mouse brain may perform fast-local and slow-global computations in parallel.

**AUTHOR SUMMARY:** In this study we analysed publicly available neural population electrophysiology data from nine brain regions in awake mice. To discover neural ensembles, we applied community detection algorithms to the spike count correlation matrices estimated from the data. We repeated the analysis at different timescales, ranging from 10 milliseconds to 3 seconds. We found that at fast timescales *<* 1 s, neural ensembles tended to be localised within single brain regions. In contrast at slower timescales of *>* 1 s, ensembles tended to be include neurons spread across multiple brain regions. Most of this effect was due to high-firing-rate neurons.

## INTRODUCTION

The brain is traditionally parcellated into anatomical regions that perform distinct computations (Zilles & Amunts, 2010). However these regions do not operate independently: successful brain function must also involve computations spread over multiple regions (Bassett & Bullmore, 2009; Power et al., 2011; Sporns & Betzel, 2016). It is unclear how local computations within a single brain region are coordinated with global computations spread across many brain regions. Several possibilities have been proposed: synchronous oscillatory activity may bind together spatially separated neural signals (Berger, Warren, Normann, Arieli, & Grün, 2007; Engel, Schölvinck, & Lewis, 2021; Fries, 2015; Gray, König, Engel, & Singer, 1989; Siegel, Donner, & Engel, 2012); travelling waves may propagate signals across the cortex (Muller, Chavane, Reynolds, & Sejnowski, 2018); or a hierarchy of timescales may separate low-level sensory processing from higher-level cognitive computations in the brain (Murray et al., 2014; Siegle et al., 2021; Zeraati et al., 2021).

Here we tested the hypothesis that computations are local to single brain regions at fast timescales, but spread across multiple regions at slower timescales.

## RESULTS

### Spatial extent of neural correlations varies with timescale

We first characterised the magnitudes of within- and between-region neural **spike count correlations** by analysing previously published data from ∼500 neurons recorded simultaneously across 9 brain regions (frontal, sensorimotor, visual, and retrosplenial cortex, hippocampus, striatum, thalamus, and midbrain) in awake mice Steinmetz, Pachitariu, Stringer, Carandini, and Harris (2019); Stringer et al. (2019) (Figure 1a,b). We calculated spike count correlations for each pair of neurons in the dataset over a range of different time bin widths, from 10 milliseconds to 3 seconds.

**Figure 1.**
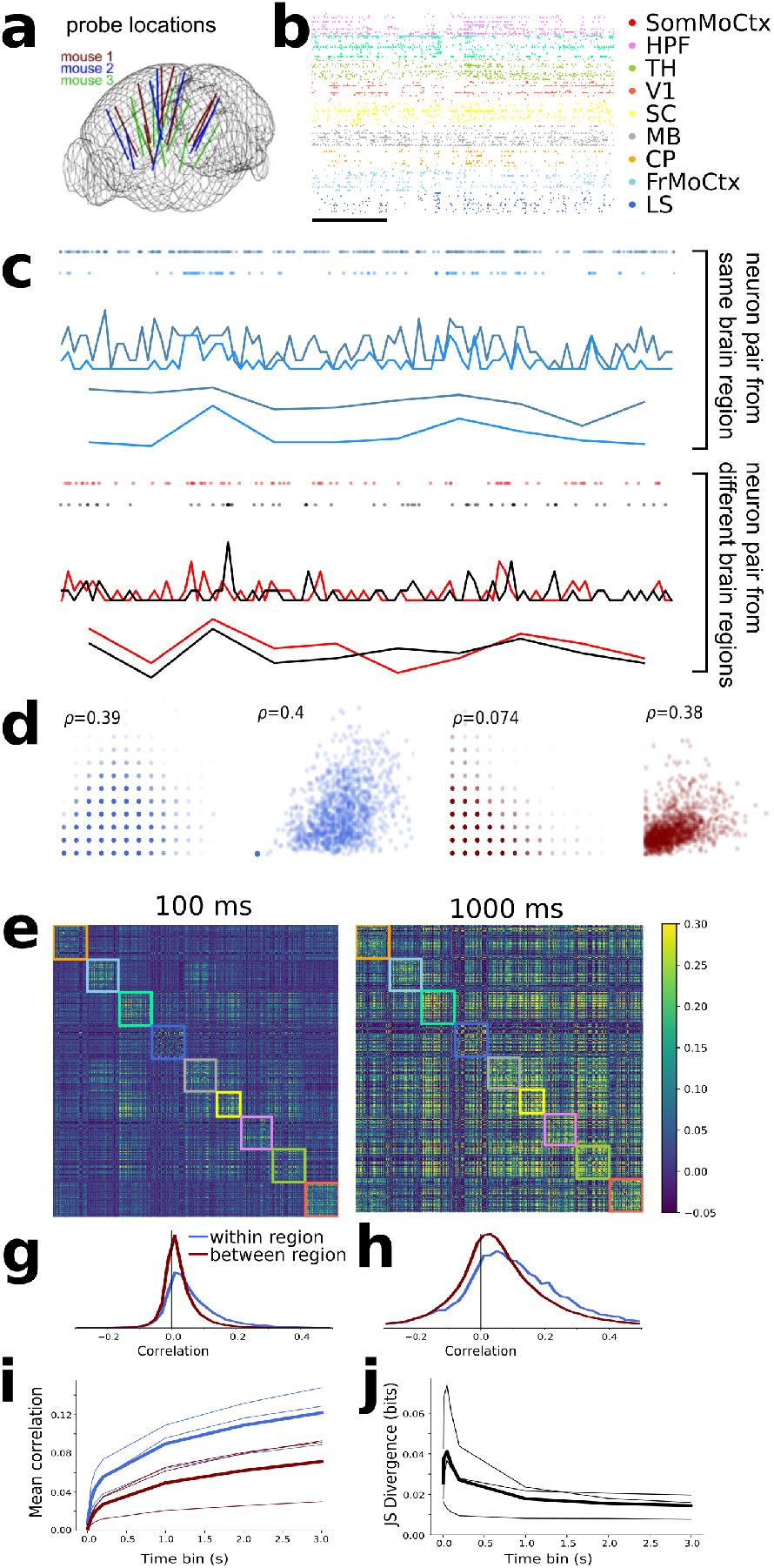
Within- and between-region neural correlations are more similar at slow timescales than fast timescales. **a** Neuropixels probe locations in the three mouse brains (adapted from Stringer et al. (2019)). **b:** Raster plot of spikes from 198 sample units from one mouse. Scale bar corresponds to 1 s. **c:** Spike-count time series from a pair of neurons recorded in the same brain region (top) and pair recorded from different regions (bottom). **d:** Scatter plots of all spike counts for same pairs of neurons shown in panel c. The x-axis shows spike count from one neuron, y-axis spike counts from the other neuron. From left to right: within region pair at 100 ms and 1000 ms time bins; different-region pair at 100 ms and 1000 ms time bins. Inset indicates pairwise spike-count correlation coefficient. **e, f:** Correlation matrix for spike counts from 494 neurons recorded from one animal with a time bin of 100 ms (e) or 1 second (f). **g, h:** Histograms of pairwise correlations from matrices in c and d for within- and between-region pairs of neurons (colours blue and red respectively) for 100 ms (g) or 1 second (h) time bins. **i:** Mean pairwise correlations as a function of time bin. **j:** Jensen-Shannon divergence of within vs between-region correlation distributions as a function of time bin.

Figure 1c shows example 10-second raster plots and corresponding spike count time series from a pair of neurons within the same brain region (light and dark blue, top) and a pair of neurons from two different brain regions (red and black, bottom) for both 100 ms and 1 s time bins. We choose these two timescales as representative examples throughout the paper. Figure 1d shows scatter plots of the spike counts for the same neuron pairs. The within-region cell pair showed the same high spike count correlation of *ρ* ≈ 0.4 at both 100 ms and 1 s time bins. In contrast, the between-region pair showed a low spike count correlation of 0.07 at fast 100 ms time bins, but a high correlation of 0.4 at slower 1 second time bins.

This general pattern held up across the dataset: Figure 1e shows the pairwise correlation matrices for all 494 neurons analysed from this animal for both the 100 ms and 1 second time bin sizes. The rows and columns of these matrices are ordered by brain region, so within-region correlations are inside the coloured boxes along the main diagonal (each colour represents a different brain region). With 100 ms bins, the within-region correlations appear stronger than the between-region correlations. However with 1 second time bins, the within- and between-region correlations appear visually similar. To explore this phenomenon, we separately histogrammed the within- and between-region values from the correlation matrices (Figure 1g,h). Both the mean (Figure 1i) and the width of correlation histograms increased with time bin size, for both within- and between-region correlations (Bair, Zohary, & Newsome, 2001).

However, the within-region correlations had a heavier positive tail than the between-region correlations at fast timescales, but markedly less so at slow timescales (Figure 1g,h). To quantify this effect, we calculated the Jensen-Shannon (JS) divergence between the two distributions. High divergence values imply greater differences in the distributions. Indeed the JS divergence decreased as a function of time bin size, consistently for the data from all three animals (Figure 1j). These results imply that at fast timescales, correlations are high only between neurons within brain regions, but at slow timescales within- and between-region neural correlations are similar.

### Low firing rate neurons preferentially correlate within brain region

Low- and high-firing rate neurons have previously been shown to serve different functions in neural circuits (Gava et al., 2021; Levenstein, Watson, Rinzel, & Buzsaáki, 2017). To test whether this dissociation is also visible in the within-vs between-region correlation structure, we plotted correlation values against geometric mean firing rate for each pair of neurons in the dataset (Figure 2a–d). Most pairs of neurons had geometric mean firing rates between 1–10 Hz (Figure 2e). Correlations tended to get stronger as a function of firing rate, for both within- and between-region pairs (Figure 2a–d)

**Figure 2.**
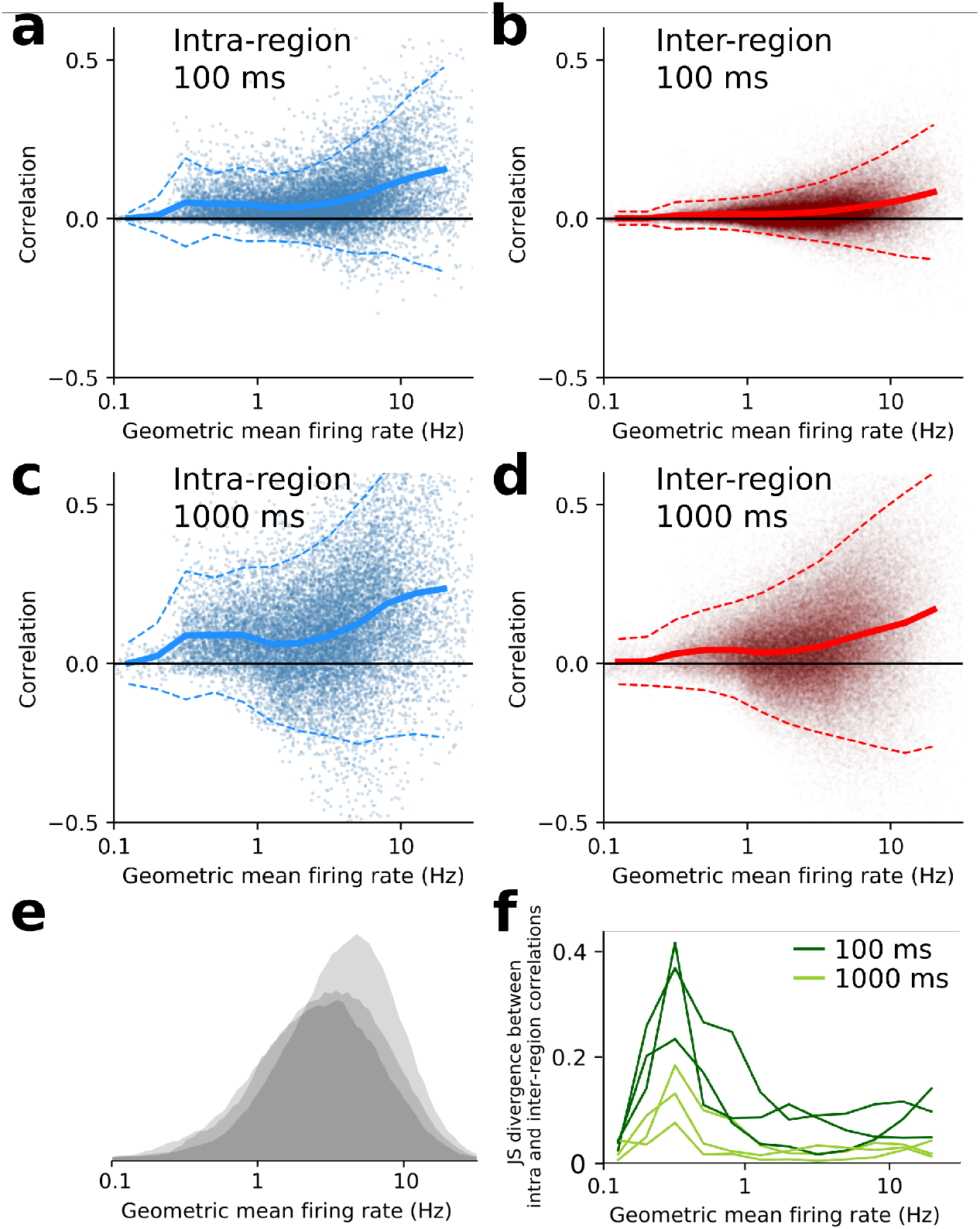
Low firing rate neurons preferentially correlate within brain regions. **a–d:** Pairwise neural correlations vs geometric mean of firing rate for many pairs from one animal, for intra (a,c) and inter (b,d) region neuron pairs, with time bin interval shown in panel insets. Solid line shows mean correlation, dashed lines are *±* 2 s.d. from mean. **e:** Histograms of all pairwise geometric mean firing rates for all three animals. **f:** Jensen-Shannon divergence between intra and inter-region correlations as a function of geometric mean firing rate, for all three animals. Dark green corresponds to spikes binned at 100 ms intervals, light green is 1000 ms intervals.

(De La Rocha, Doiron, Shea-Brown, Josicá, & Reyes, 2007). We binned pairs by their geometric mean firing rate and calculated the JS divergence between the within- and between-region correlations as a function of firing rate bin (Figure 2f). At both fast and slow timescales, low-firing rate pairs had stronger within-region correlations than between-region correlations. In contrast, high firing rate pairs had moderate divergence at 100 ms timebins and almost zero divergence at 1 second time bins. This implies that high-firing rate neurons correlate almost equally strongly within- and between-regions, but low-firing rate pairs have similarly low within- and between-region correlations at all timescales. Therefore the phenomenon seen in Figure 1 is mainly due to high-firing rate neuron pairs.

### Detected ensembles align with anatomical regions at short time bins, but not long time bins

To test if **neural ensembles** also showed different structure at fast and slow timescales, we ran a community detection algorithm from network science on the correlation matrices to detect ensembles (Figure 3a) (Humphries, Caballero, Evans, Maggi, & Singh, 2021). The algorithm splits the neurons into non-overlapping subsets based on their correlations, trying to discover ensembles of neurons with strong positive correlations between the members of each ensemble, but weaker correlations with neurons in other ensembles (Methods). Figure 3b and c shows the same example correlation matrices from Figure 1e, but with the rows and columns reordered by ensemble membership. In all three animals we found fewer ensembles at longer time bin sizes (Figure 3f). Crucially, the ensemble detection algorithm did not know anything about which brain regions each neuron belonged to. To visualise the brain region membership of each ensemble, we plotted a small square for each neuron coloured according to its brain region (Figure 3d,e). At 100 ms time bins, most ensembles contained neurons from only a small number of brain regions, whereas at 1 second time bins almost all ensembles contained neurons from several brain regions. To quantify this effect, we asked the questions: what is the probability that any arbitrary neuron pair is in the same ensemble? And does this differ for pairs of neurons within the same brain region vs pairs across two brain regions? 20–30% of same-region pairs were in the same ensemble, but only 10–20% of different-region pairs were in the same ensemble (Figure 3g). The difference between these two fractions decreased towards zero as a function of time bin size (Figure 3h), implying that at fast time scales neurons in the same brain region had a higher chance of being in the same ensemble than two neurons in different brain regions, but this distinction got weaker at slower time scales. To further quantify the effect, we used a distance measure from information theory to ask the question: how different are the sets of neuron groups when defined by brain region vs defined by ensemble? (Methods: clustering comparison). This ‘variation of information’ measure increased as a function of time bin size in all three animals (Figure 3i), again implying that anatomical regions and neural activity ensembles are more similar at fast timescales than slow timescales.

**Figure 3.**
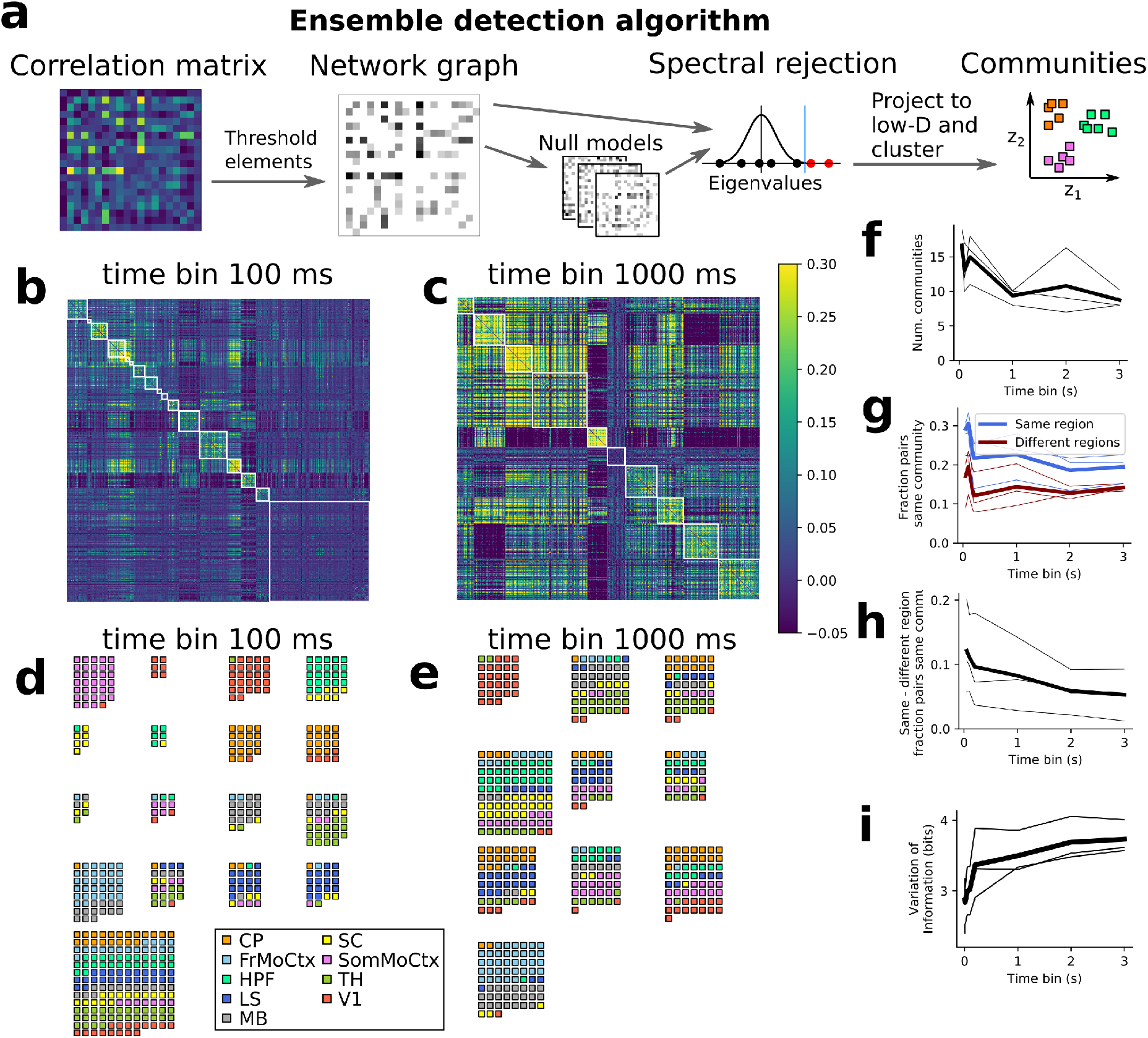
Neural ensembles are within-region at fast timescales but multi-region at slow timescales. **a:** Schematic diagram of community detection algorithm steps. **b, c:** Same correlation matrices as fig 1 sorted by ensemble. **d, e:** Example ensembles at short (d) and long (e) timescales. **f:** Number of detected ensembles vs time bin size. **g:** Fraction of same and different region neuron pairs being in same community, vs time bin size. **h:** Difference in fraction of same and different region neuron pairs being in same community (same data as panel g). **i:** Variation of information (measure of dissimilarity of anatomical vs ensemble partitions) vs time bin size.

## DISCUSSION

Although previous studies have compared within- and between-region neural correlations, to our knowledge none have described the fast-local vs slow-global ensembles phenomenon we presented here. There are a few possible reasons for this gap: most electrophysiological studies either looked at small numbers of neurons where the phenomenon may not be statistically detectable, or looked at aggregate neural activity measures such as local field potentials (Engel et al., 2021) which would miss the single-neuron-resolution ensembles we discovered. Modern large-scale two-photon imaging methods do enable simultaneous recordings from single neurons in multiple brain regions, but with poorer signal-to-noise and slow sampling rate so also may not be able detect the phenomenon we described. Non-invasive methods for recording brain activity such as functional Magnetic Resonance Imaging, magneto- and electro-encephalography do not have high enough spatial resolution to isolate single neurons, so will not be able to resolve these ensembles which are often spatially intermingled.

We examined this phenomenon only for 9 particular brain regions, which despite all exhibiting the effect, differed in their mean firing rates and correlations (Stringer et al., 2019), and presumably also differ in the computations that they perform for the brain at large. It would be interesting to try to understand if and how each brain region adapts variations of the general fast-local, slow-global principle and relate it to its overall function. For example, it may be the case that regions pairs which have strong, direct anatomical connections between them (Oh et al., 2014) show stronger correlations, or prefer to correlate on fast timescales.

There are several important limitations to our analysis. First, we used Pearson’s correlation as our measure of co-activity between neurons. This captures only linear dependencies so may miss nonlinear interactions, but crucially is also averaged over the entire 1-hour recordings. Therefore it may miss transient co-activity events such as ripple oscillations, which can synchronize neural activity across multiple brain regions (Jadhav, Rothschild, Roumis, & Frank, 2016). Second, we defined ensembles as groups of neurons with positively correlated activity, which ignores the fact that negative correlations may also be indicative of an interaction between neurons. However we found qualitatively consistent results when we ran our ensemble detection algorithm on graphs defined by absolute values of the correlations. In contrast, running the algorithm based on only negative-correlation interactions did not reproduce the fast-local vs slow global effect. Third, we assumed for simplicity that ensembles were non-overlapping. However the correlation matrices in Figure 3b,c show substantial structure outside the detected communities, which implies that at least some neurons participate in multiple ensembles. It would be interesting for future studies to explore the fast-local, slow-global phenomenon in overlapping neuron groups using alternative community-detection algorithms (Xie, Kelley, & Szymanski, 2013).

Fourth, we examined neural co-activity only in the time range of 10 ms to 3 seconds, but there may well be important co-activity dynamics at both faster and slower timescales. For fast timescales, 1–10 ms, spike count correlations would not be an appropriate measure because most neurons in this dataset fired at *<* 10 Hz (Figure 2e), so almost all time bins would have zero spike counts and correlations would be very close to zero (Figure 1i). Future studies exploring fast timescales could use alternative measures, such as spike time cross-correlograms (Berger et al., 2007; Cohen & Kohn, 2011). Conversely, there may also be interesting cross-brain neural ensemble dynamics at the ‘infra-slow’ timescales ∼10–100 s, which has previously been shown to have qualitatively different features to neural population dynamics at fast timescales (Okun, Steinmetz, Lak, Dervinis, & Harris, 2019).

It is important to note that these results do not imply that all local computations are fast while all global computations are slow. Indeed Figure 3d shows some evidence for mixed-region ensembles at fast timescales. There are also established counter-examples to the phenomenon, such as *>*1 s cell-intrinsic persistent spiking activity (Egorov, Hamam, Franseán, Hasselmo, & Alonso, 2002; Lüthi & McCormick,1999), and *>* 100 Hz fast ripple oscillations that synchronize distal brain regions (Jadhav et al., 2016). We also did not explore how these effects vary with the behavioural state of the animal, or the sensory stimuli it is exposed to. For example, behavioural state fluctuations have widespread effects on neural activity across the brain at *>*1 s timescales (Stringer et al., 2019). Therefore it may be that behavioral state changes are a major contributor to the slow-global co-ordination effects we report – future studies could assess this by manipulating animal behavior or by statistically regressing out the effect of behaviour measures on neural co-activity. In summary, we consider the fast-local, slow-global phenomenon we describe here as only an average tendency for neural activity co-ordination in the mouse brain, which may act as a scaffold for other spatiotemporal dynamical structures to rest upon.

Why might the fast-local vs slow-global dissociation exist? From a mechanistic point of view, one explanation may be that the energetic and space constraints on brain wiring imply that long-range, between-region signals can be transmitted only at low-bandwidth and with some latency (Sterling & Laughlin, 2015). There are typically fewer long-range synaptic connections than local connections, between-region signalling is low-dimensional (Semedo, Zandvakili, Machens, Byron, & Kohn, 2019), and mammalian axons transmit action potentials between brain regions with latencies of 10–100 ms (Swadlow & Waxman, 2012). These bandwidth and latency constraints will limit the speed of any computations that require back-and-forth recurrent signalling between neurons. This issue is well known in human-made computers, where the ‘von Neumann bottleneck’ for transferring data between memory and CPU via low-bandwidth and high-latency databuses constrains computation speed (Hennessy & Patterson, 2011). From a functional point of view, a separation of timescales between local and global computations may allow for less interference between processes, and allow local neural circuits to complete their tasks quickly before broadcasting the results to other brain regions (Engel et al., 2021).

## METHODS

### Original data description

All data analysed in this study were sourced from a publicly available dataset (Steinmetz et al., 2019). The experimental procedures have been described previously (Stringer et al., 2019). Briefly, eight Neuropixel probes were used to record electrophysiological activity simultaneously from nine brain areas: frontal, sensorimotor, visual, and retrosplenial cortex, hippocampus, striatum, thalamus, and midbrain, in each of three 10–16 week-old mice. The mice were awake but head-fixed. 2296, 2668, and 1462 stable units were isolated from each mouse, respectively, across a ∼1-hour recording. Spikes were sorted using the Kilosort2 algorithm (Stringer et al., 2019). The published dataset lists spike times for each of the units.

### Code availability

All analyses were performed using Python, and figures were prepared using Inkscape. The computer code we used for calculating neural correlations is available at: https://github.com/thomasjdelaney/Regional_Correlations.

We also implemented the community detection algorithm from Humphries et al. (2021) in our own Python code:

https://github.com/thomasjdelaney/Network_Noise_Rejection_Python. Code for calculating the firing rate dependencies of correlations and for generating the figures is available at: https://github.com/odonnellgroup/fast-local-slow-global-ensembles.

### Conversion to spike count data

We selected a random subset of ∼500 neurons for each mouse, approximately balancing the number of neurons from each of the nine brain regions so as not to introduces regional biases into our analyses. We transformed the spike timing data into binned spike count data by dividing the experimental period into time bins, and counting the spikes fired by each unit per time bin. We varied the time bin size from 10 ms to 3000 ms.

### Correlation coefficients

We used the python function `scipy.stats.pearsonr` to calculate Pearson’s sample correlation coefficient for spike counts from each pair of neurons:

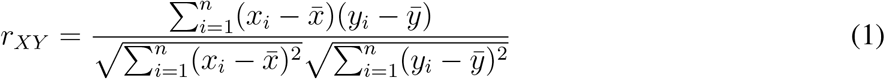

where {(*x*_*i*_, *y*_*i*_)} for *i* ∈ {1, …, *n*} are the paired samples from neurons *X* and *Y*, and 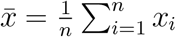, and 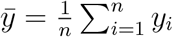 are the sample means.

### Ensemble detection analysis

The correlation matrices can be interpreted as an undirected weighted graph between the neurons, where the weight of each connection is equal to the correlation between each pair of neurons.

The community detection algorithm we used to detect neural ensembles (Humphries et al., 2021) is designed for networks with positively weighted connections, but many neuron pairs were negatively correlated. To adapt the correlation matrices for the algorithm we rectified the network by setting all the negative weights to zero.

We also wanted to exclude any correlations that could be judged to exist ‘by chance’. To do this, for each neuron pair and time bin width we shuffled the spike count time series, calculated correlations from many shuffle permutations, then found the 5th and 95th percentiles of the shuffled correlation distributions. We used these percentiles as detection thresholds for the data correlations, and set any intermediate correlation values to 0. This excluded any ‘chance’ correlations from our network and created a sparser network.

We now give an overview the community detection algorithm. Full details are available in Humphries et al. (2021). Given some network represented by an adjacency matrix **A**, a community within that network is defined as a collection of nodes where the number of connections within these nodes is higher than the expected number of connections between these nodes. In order to quantify the ‘expected’ number of connections, we need a model of random networks with little or no structure, analogous to a ‘null model’ in traditional hypothesis testing. Since we are working with weighted sparse networks, we used a *weighted configuration model*, a canonical null network model for weighted networks. This model preserves the degree sequence and weight sequence of each node in the data network, but with the edges distributed randomly (Fosdick, Larremore, Nishimura, & Ugander, 2018). Any structure in the data-derived network beyond its degree sequence and weight sequence will not be captured by the weighted configuration model. In practice we used an extension that also preserves sparsity, by sampling from a probability distribution for the creation or non-creation of each possible connection, then distributing the weight of the data network randomly in this sparse network (Humphries et al., 2021). To detect the structure in the data beyond that seen in the null model we used a spectral rejection procedure, as follows.

Given a data network matrix **W** and expected network of our null network model ⟨**P**⟩, then the departure of our data network from the null network can be described by the ‘deviation matrix’

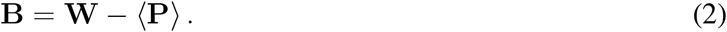

A common choice for ⟨**P**⟩ in community detection is the ‘configuration model’ (Fosdick et al., 2018; Humphries et al., 2021).

To test for structure in the network **W**, we examine the eigenspectrum of **B** and compare it to the eigenspectrum of the null model. Since our data model doesn’t allow self loops and is not directed, the matrix representing the network will be symmetric and positive semi-definite, and will therefore be invertible with real eigenvalues. We selected a null model with the same characteristics.

To find the eigenspectrum of the null model, we generated *N* samples from our null model *P*_1_, …, *P*_*N*_, and we measured their deviation matrices *B*_1_, …, *B*_*N*_. We then calculated the eigenspectrum of each of those samples. We calculated the upper bound of the null model eigenspectrum by taking the mean of the largest eigenvalues of *B*_1_, …, *B*_*N*_ .

We then calculated the eigenspectrum of **B**, our data network deviation matrix. Eigenvalues above the upper bound of the null model eigenspectrum give evidence for community structure in the data network. If there are *d* data eigenvalues lying outside of the null network eigenspectrum, the *d* eigenvectors corresponding to these eigenvalues form a vector space. If we project the nodes of our network into this vector space, by projecting either rows or columns of the data matrix, we can see how strongly each node contributes to the vector space. Nodes that contribute strongly to the additional structure will project far away from the origin, nodes that do not contribute to the additional structure will project close to the origin. We used this information to discard those nodes that do not contribute.

To detect the neural ensembles we first project all of the nodes into this *d*-dimensional subspace, then perform the clustering. The clustering and community detection procedure is described in (Humphries et al., 2021).

In practice, the procedure is carried out *n* times (we chose *n* = 100 times), this returns *n* clusterings. We resolve these *n* clusterings to one final clustering using consensus clustering. We used the consensus clustering method that uses an explicit null model for the consensus matrix, as outlined in (Humphries et al., 2021).

### Clustering Comparison

A clustering C is a partition of a set *D* into sets *C*_1_, *C*_2_, …, *C*_*K*_, called clusters, that satisfy the following for all *k, l* ∈ {1, …, *K*}:

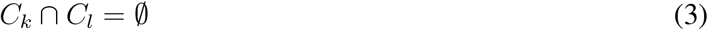

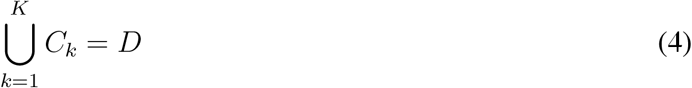

If we consider two clusterings, C with clusters *C*_1_, *C*_2_, …, *C*_*K*_ and C_*1*_ with clusters 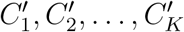. There are a number of possible measurements to compare C and C_*1*_, we used the ‘variation of information’ (VI). This is an information theoretical quantity based on the mutual information, defined as

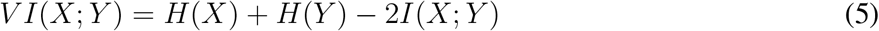

We can rewrite this as the summation of two positive quantities

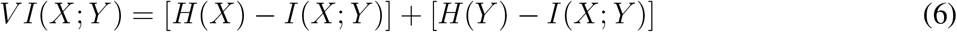

It is the summation of the uncertainty in the random variables *X* and *Y* excluding the uncertainty shared by those variables. It forms a metric on the space of clusterings. That is,

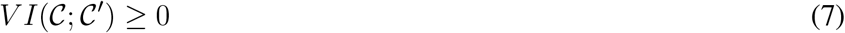

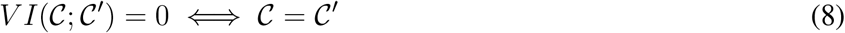

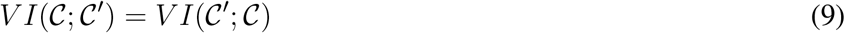

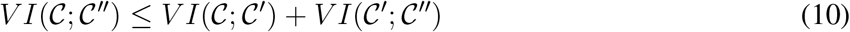

In order to quantify the difference or similarity between the communities detected in our correlation network and the anatomical classification of the cells in that network, we considered the communities and the anatomical regions as two different clusterings, 𝒞_*comm*_ and 𝒞_*anat*_, respectively. We measured the distance between the clusterings using the variation of information. This quantity is zero if the two clusterings are identical and large if the clusterings are dissimilar. We calculated this quantity using custom Python code.

## ACKNOWLEDGMENTS

We thank Michael Ashby for useful discussions and Nick Steinmetz for making the data public. Funding was provided by the EPSRC (Doctoral Training Partnership studentship to TJD) and MRC (MR/S026630/1 to COD).

## TECHNICAL TERMS

**Neural ensemble:** a group of neurons that are often active at the same time, likely participating in a joint representation or computation.

**Spike count correlations:** neurons communicate by emitting action potentials, or spikes. The rate of these spikes tend to be correlated between pairs of neurons. Spike count correlation is a statistical measure of the strength of this correlation.

**Community detection algorithm** : an algorithm which tries to split items into groups such that the connections between items in a group tend to be stronger than for items in different groups.

**Low-dimensional:** the idea that the signal carried by a large number of items can be compressed and represented by a smaller number of factors without much loss of information.

